# Closed-Loop Control of Active Sensing Movements Regulates Sensory Slip

**DOI:** 10.1101/366609

**Authors:** Debojyoti Biswas, Luke A. Arend, Sarah A. Stamper, Balázs P. Vágvölgyi, Eric S. Fortune, Noah J. Cowan

**Affiliations:** Department of Electrical and Computer Engineering, Johns Hopkins University, 3400 N. Charles Street, Baltimore, MD 21218, USA; Laboratory for Computational Sensing and Robotics, Johns Hopkins University, 3400 N. Charles Street, Baltimore, MD 21218, USA; Department of Mechanical Engineering, Johns Hopkins University, 3400 N. Charles Street, Baltimore, MD 21218, USA; Federated Department of Biological Sciences, New Jersey Institute of Technology, 323 Dr. Martin Luther King Jr. Bulevard, Newark, NJ 07102, USA

**Keywords:** active sensing, reafferent feedback, control theory

## Abstract

Active sensing involves the production of motor signals for the purpose of acquiring sensory information [1–3]. The most common form of active sensing, found across animal taxa and behaviors, involves the generation of movements—e.g. whisking [4–6], touching [7,8], sniffing [9,10], and eye movements [11]. Active-sensing movements profoundly affect the information carried by sensory feedback pathways [12–15] and are modulated by both top-down goals (e.g. measuring weight vs. texture [1,16]) and bottom-up stimuli (e.g. lights on/off [12]) but it remains unclear if and how these movements are controlled in relation to the ongoing feedback they generate. To investigate the control of movements for active sensing, we created an experimental apparatus for freely swimming weakly electric fish, *Eigenmannia virescens*, that modulates the gain of reafferent feedback by adjusting the position of a refuge based on real time videographic measurements of fish position. We discovered that fish robustly regulate sensory slip via closed-loop control of active-sensing movements. Specifically, as fish performed the task of maintaining position inside the refuge [17–22], they dramatically up- or down-regulated fore-aft active-sensing movements in relation to a 4-fold change of experimentally modulated reafferent gain. These changes in swimming movements served to maintain a constant magnitude of sensory slip. The magnitude of sensory slip depended on the presence or absence of visual cues. These results indicate that fish use two controllers: one that controls the acquisition of information by regulating feedback from active sensing movements, and another that maintains position in the refuge, a control structure that may be ubiquitous in animals [23,24].

## Results

### Active sensing is modulated by reafferent gain

In active sensing, an animal stimulates and/or regulates the information available to its own sensory systems via movement or, in a handful of specialized animals, via the generation of sensory signals such as electric fields or echolocation calls [25–28]. A hallmark of active sensing is that it is modulated in relation to changes in behavioral or sensory context. For example, as the weakly electric fish *Eigenmannia virescens* maintains its position within a refuge, it also produces small fore-aft body movements for active sensing [12]. These movements create a dynamic difference between the position of the fish and the refuge, i.e. a *sensory slip* analogous to retinal slip [23,29], albeit mediated by the propagation of electricity in water [28]. Active swimming movements in electric fishes likely prevent perceptual fading and enhance spatiotemporal patterns of sensory feedback [12,13,30] serving a similar role as small eye movements in vision [15,30,31].

Here we address whether active movements depend on the electrosensory feedback they produce (i.e., are the active movements under closed-loop control) or whether active movements are independent of this feedback (i.e., open loop). To examine this, we designed a system in which the sensory slip experienced by the animal can be artificially augmented by an experimentally defined gain. In this system, custom software tracks the position of the fish and adjusts the position of the refuge in real time (Fig. 1 A,B); the movement of the refuge is determined by the movement of the fish based on an experimental gain, γ (Fig. 1 C).

**Figure 1:**
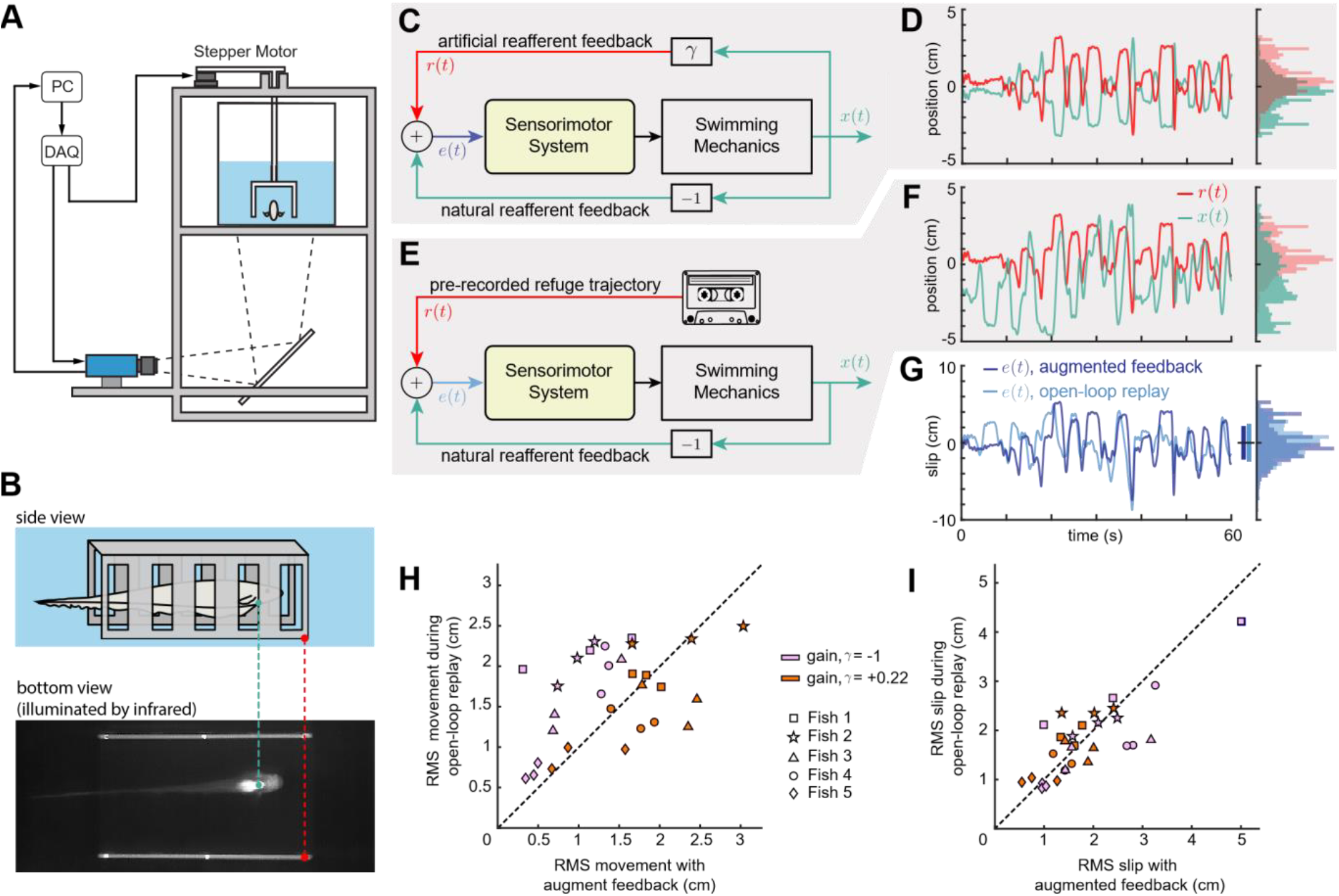
Categorical differences between augmented feedback experiments and open-loop replay experiments. (A) Schematic showing a front view *E. virescens* inside a moving refuge. The refuge is actuated by a stepper motor, controlled by a computer that processes video images streamed from a camera in real time. The camera is positioned to observe the fish and refuge from below via a mirror. (B) Side view (schematic) and bottom view (image) of a fish inside the refuge. The fish has a ventral bright patch (aqua dot) in the infrared-illuminated video that can be tracked as the fish swims inside the refuge (red dot). (C) Augmented feedback experiment. The fish position, *x*(*t*), is multiplied by the augmented gain γ to control the refuge position, *r*(*t*). (D) Sample data of *x(t)* and *r(t)* for an augmented feedback experiment with gain γ = −1 with the corresponding histograms; aqua is the position of the fish and red the refuge. All histograms are plotted to the same scale (E) Open-loop replay experiment. A pre-recorded refuge trajectory from the augmented feedback experiment is used as refuge motion *r*(*t*). (F) Sample data of *x(t)* and *r(t)* from a replay experiment in which the refuge trajectory, *r*(*t*) from (D) was replayed. Fish trajectories in (D) and (F) are markedly different despite identical refuge trajectories. (G) Sensory slip is plotted for both the sample data of (D) and (F). The length of the small vertical bars denotes 2×*SD* of *e*(*t*). (H) The root-mean-square (RMS) of movements differed between augmented feedback and replay conditions, and depended on γ. Each marker shape represents a different fish, each point is the mean of 5 trials. Violet indicates a gain of γ = −1 and orange γ = +0.22. The dashed line corresponds to the null hypothesis of equal RMS active movement for both gains. (I) Sensory slip between experimental conditions was maintained, irrespective of gain. Colors and marker shapes as in (H). See also Supplemental Videos.

This system permits the experimental modulation of reafferent feedback. In the veridical condition, the fish experiences an equal-but-opposite sensory slip to its own movement (reafferent gain of −1). When the refuge, *r*(*t*), is moved in relation to the fish, *x*(*t*), via an experimentally augmented gain, γ, the refuge motion is given by *r*(*t*) = γ*x*(*t*). Thus, the sensory slip, or “error”, is given as follows:

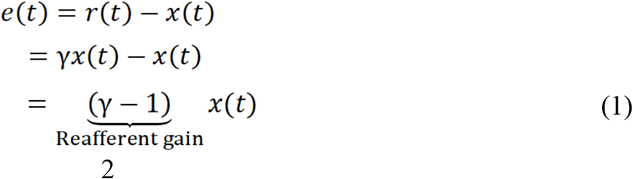

For example, when the augmented gain γ is set to 0.22, the movement of the refuge follows the position of the fish but at a smaller amplitude, thereby reducing the sensory slip (reafferent gain of −0.78). When γ = −1 the refuge moves as fast as the fish, but in the opposite direction (reafferent gain of −2; see Fig. 1 D). See Supplemental Videos. When γ = 0, the fish’s reafferent feedback is −1, the natural self-motion gain that an animal experiences when it moves.

We performed two categories of experiments, **augmented feedback experiments** and **open-loop replay experiments**. In the augmented feedback experiments, the animals performed station-keeping in the apparatus described above.

In all prior studies of refuge station-keeping in these fish [17–21], there was no augmented feedback (γ = 0, veridical) and an external refuge motion was applied, creating a time-varying exafferent signal. In our augmented feedback experiments, by contrast, the reafferent gain was experimentally manipulated but the refuge motion was otherwise held constant, i.e. motions of the refuge were driven by the fish and not by an external reference command.

In replay experiments, γ was set to 0, and we replayed trajectories of the refuge that were recorded during previous closed-loop trials (Fig. 1 E,F). The movement of the refuge was identical across open- and closed-loop trials making the key difference between these trials whether or not a closed-loop coupling existed between the movements of the fish and those of the refuge. This manipulation isolated the effects of enhancing or suppressing sensory feedback from the details of the refuge trajectory, since the latter is identical across open- and closed-loop experiments.

As a measure of animal’s movement and the resulting sensory slip, we calculated the root-mean-squared (RMS) of both signals (with means subtracted), a common measure of signal magnitude:

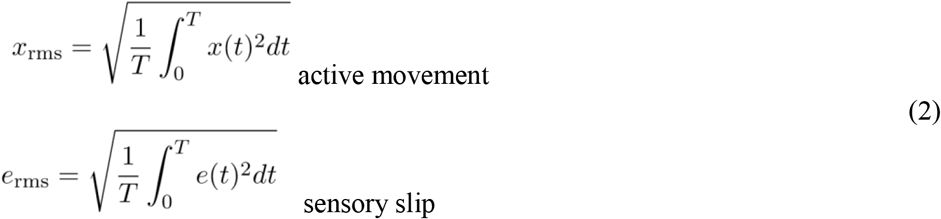

If active movements did not rely on feedback control, these movements would be similar across open- and closed-loop trials. However, we observed less movement in closed-loop trials with a negative gain (enhanced reafference) than during the corresponding open-loop replay trials (Fig. 1 H). Conversely, with a positive gain (attenuated reafference) gain, we observed more movement in closed-loop trials than in the corresponding open-loop trials. The ratio of open-loop to closed-loop RMS movements were significantly different depending on the augmented gain (for γ = −1: ratio = 2.033, SEM = 0.31; for γ = +0.22: ratio = 0.912, SEM = 0.06, across fish; Mann-Whitney-Wilcoxon test: *p* = 0.008 (two-tailed)).

In contrast, we observed no significant difference between RMS sensory slip in closed-loop and open-loop replay trials (Fig. 1 G,I; for γ = −1: ratio = 0.963, SEM = 0.11; for γ = +0.22: ratio = 1.147, SEM = 0.08, across fish; Mann-Whitney-Wilcoxon test: *p*=0.310 (two-tailed)).

Taken together, these results demonstrate that reafferent feedback is used to control movements for active sensing. The question remains, what is the goal of this feedback control system?

### Fish regulate sensory slip across reafferent gain

Based on the theory that fore-aft movements are for the purpose of sensing, we hypothesized that such movements would be regulated based on ongoing reafferent feedback. Specifically, our hypothesis is that fish maintain sensory slip, i.e. fish will swim more or less as needed to maintain a constant RMS positional slip. Intuitively, if reafferent feedback is enhanced, fish should swim less, and, if reafferent feedback is suppressed, fish should swim further. We experimentally varied the reafferent gain, γ−1, over the range −2 to −0.5 and measured the resulting RMS sensory slip. Previous work has shown that active-sensing movements differ dramatically depending on lighting: in the dark, the movements are much larger than in the light [12] so we investigated a second hypothesis that the set point for this regulation of sensory slip depends on the lighting condition.

From Eqn. (1), the sensory slip experienced by the fish in the augmented feedback apparatus with augmented gain γ is given as follows:

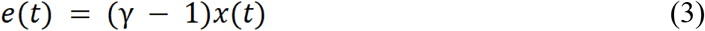

Given this relation, a straight-forward calculation shows that the root-mean-square of the slip signal, *e*_rms_, and the fish position, *x*_rms_, for each gain and lighting condition must be related by a constant factor:

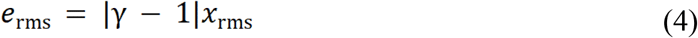

We selected gains that ensure 1 − γ > 0, and hence:

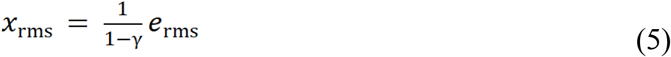

Defining transformed gain, β=(1−γ)^−1^ simplifies this equation:

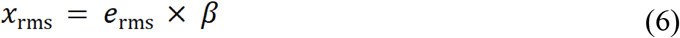

Therefore, if the fish were to maintain constant RMS slip *e_rms_*(invariant to the reafferent gain), then *x*_rms_ would be a linear function of the transformed gain β, with zero intercept and with slope given by the RMS slip, *erms*. More generally, if the fish were to modulate its swimming behavior to regulate RMS slip, we would expect the fish movement to increase as a function of increasing β:

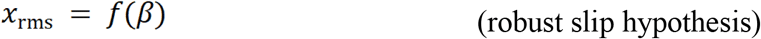

where *f*(⋅) is a monotonically increasing function.

In the null hypothesis, the active movements do not depend on the immediate feedback they create (open-loop strategy), thus we would expect the swimming motions to be invariant to applied gain:

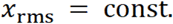

To test these hypotheses, we selected values for β to be uniformly spaced between 0.5 and 2, corresponding to γ being nonuniformly spaced between −1 and 0.5. We found that RMS movement increased as a function of β for each individual animal (Fig. 2 A,C,E; Mann-Kendall test, *p*<0.001 with positive Sen’s slope for each of *N*=6 fish for each lighting condition). Fitting a line through the origin *x_rms_*=*const*×β explained 88% of the variance across fish in the light and 65% of the variance in the dark (see Table S1; Recall that *R*^2^ is by definition zero for the null hypothesis since the constant is simply the mean.) A quadratic curve significantly improved the fit for the dark trials (adjusted *R*^2^=0.88; polynomials above order 2 did not improve adjusted *R*^2^; see Table S1 for details). The monotonically increasing RMS movements served to generate nearly constant RMS slip (Fig. 2 B,D,F). The deviation from the linear trend at high positive gains in the dark may be a consequence of a motor/energetic trade-off: the highest values of gain place an extreme demand on the animal’s locomotor system, requiring 2× more RMS movements to maintain the same RMS slip. This deviation caused the RMS slip to decrease slightly at the highest gains tested, as shown. Obviously, the locomotor limitations of the animal demands that this curve asymptote to zero as β→∞ (no reafferent gain). It is surprising, therefore, that the RMS slip is relatively flat over the large range of gains tested.

**Figure 2:**
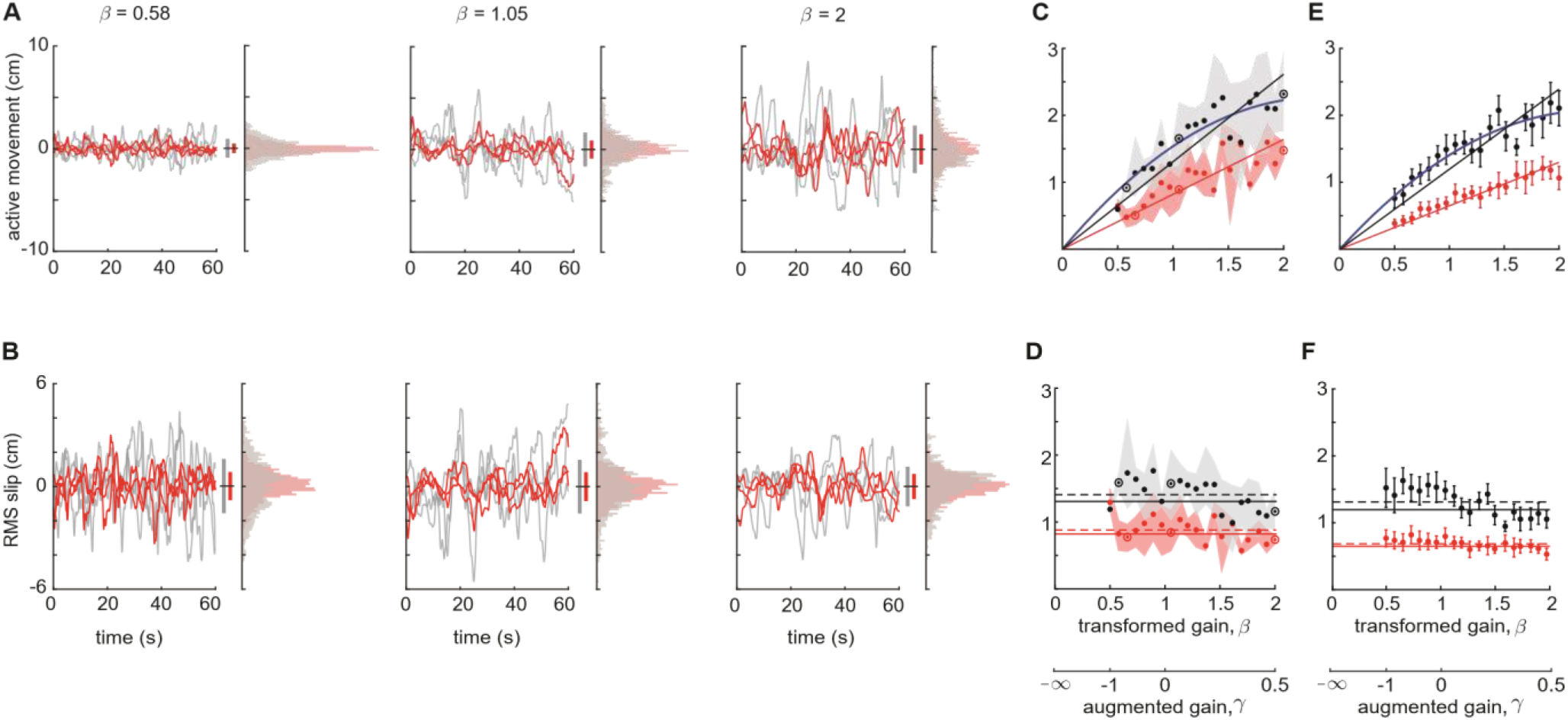
Sensory slip is maintained across reafferent gains in augmented feedback experiments. (A) Representative trajectories (mean subtracted) of one fish along with the corresponding histograms for different values of transformed gain, β (gray: ‘lights-off’, red: ‘lights-on’). Histograms approximate the probability density of *x(t)* based on the combined data from all three replicates. All histograms are plotted to the same scale. The length of the small vertical bars denotes 2×*SD* of the corresponding probability densities. (B) For the same fish, sensory slip data are presented. Format as in (A). (C) Representative data for the same fish showing that RMS movements increased as a function of transformed gain, β. Each marker represents the mean across three replicates at different gain values (black: ‘lights-off’, red: ‘lights-on’). The gain values used in (A,B) are presented as unfilled circles with dots. Shaded regions denote the maximum and minimum experimental values at each gain. Solid lines correspond to the best linear fits (through the origin, see text). A quadratic (blue, solid) improved the fit for the ‘lights-off’ trials. (D) For the same fish as in (C), RMS slip was approximately constant across β. Shaded regions and markers have the same format as in (C). The solid horizontal lines are the predicted average RMS slip based on the slope of the RMS active movement linear fits in (C). The dashed lines are the actual values for the average RMS slip. (E,F) Combined data for all individuals (*N* = 6) with mean ± SEM indicated. See also Table S1.

### To increase active sensing, fish swim farther, not faster

How do fish regulate RMS slip across gains? The most obvious means by which to increase the RMS of a signal would be to scale up the overall amplitude of active sensing movement by swimming faster. If fish simply had done that both the RMS of position and velocity would have scaled up by the same factor, and we would see RMS velocity follow an identical monotonic trend as position, holding RMS velocity slip constant. However, that was not the case: fish kept their RMS velocity movements nearly constant as a function of beta despite the dramatic increase in positional RMS movements. See Fig. 3A,C. Hence, the mechanism for increasing active-sensing movements was not to scale velocity. Further, the nearly constant velocity RMS implies that the RMS velocity slip should decrease in the shape of a hyperbole (STAR ⋆ Methods) which indeed is a good approximation (R^2^=0.88 in the light, R^2^=0.90 in the dark) as corroborated by our data (Fig. 3B,D).

**Figure 3:**
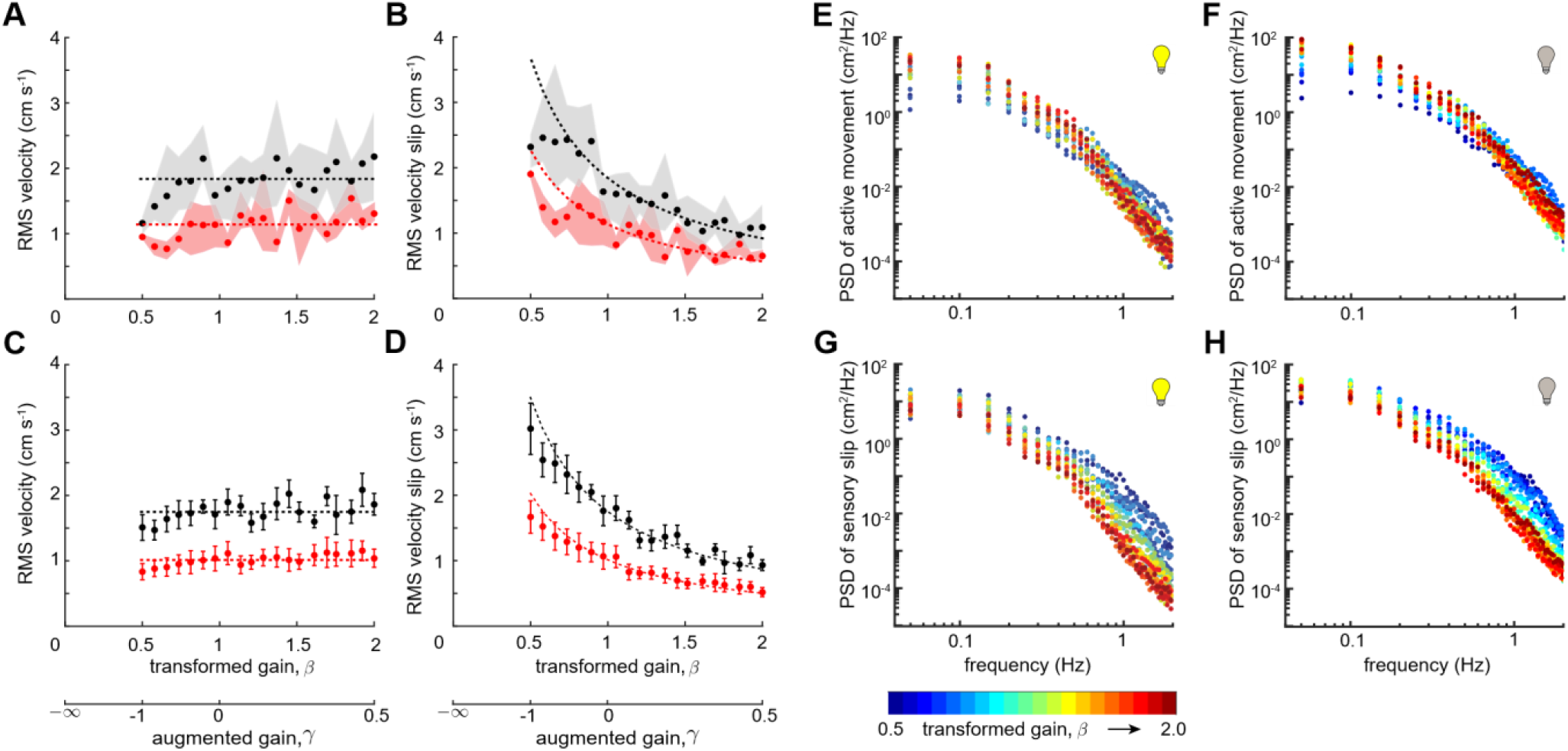
Fish do not simply increase velocity to maintain robust slip in augmented feedback experiments. (A) Representative data showing RMS velocity as a function of transformed gain, β. Each marker represents the mean across three trials and the shaded regions indicate the maximum and minimum of these trials (black: ‘lights-off’, red: ‘lights-on’). (B) The RMS velocity slip for the same fish, again averaged across three replicates. Same format as in (A). (C,D) Combined data for all individuals (*N*=6) with mean ± SEM indicated. The dashed lines in (A,C) are the constant fits to the data and in (B,D) the corresponding predictions of velocity slip. (E,F) Representative data from one fish showing PSD of fish active movements in light (E) and dark (F). For each gain (indicated by color) the PSD was averaged across three trials. At low frequencies but not high frequencies, increasing gain β is correlated with increased power (nearly 50-fold change from blue to red dots). (G,H) Representative data from the same fish showing the PSD of sensory slip (fish relative to refuge) in light (G) and in dark (H). At low frequencies, the power spectrum of slip is largely independent of gain β (less than 10-fold difference and colors overlapping). At high frequencies slip is highly gain dependent (blue dots nearly 100-fold above red). See also Figure S1,S2; Table S2,S3.

Next, we analyzed the power spectral density (PSD) of active movement and sensory slip. We observed that peaks in each fish’s active movement (Fig. 3 E,F) occur at low frequencies (< 0.1 Hz). The amplitude of these peaks increase as β was increased in both light and dark conditions (Fig. S1, Table S2). In contrast, we did not observe any overall significant change in power at higher frequencies as we changed β (in terms of −10 dB cutoff frequency, Table S2). This measure shows that fish are not relying on changes in rapid fore-aft movements, but rather use low-frequency fore-aft movements to compensate for changes in feedback gain.

How did the gain-dependent changes in the PSD of active movements affect sensory slip? The PSD of sensory slip is highest at low frequencies (Fig. 3 G,H, Fig. S1). The fish increased the low-frequency power of active movements as a function of β (Spearman’s rank correlation coefficient, *ρ*_s_ ≥ 0.71, *N*=6), thereby maintaining the low-frequency power of slip. Further, the fish did not significantly alter high-frequency power of active movements as a function of β, and therefore the power of sensory slip at high frequencies decreased with increased gain β (*ρ*_s_ ≤ −0.82, *N*=6). To quantify the transition from low to high frequency, we measured the −10 dB cutoff (0.1 amplitude crossover) frequency and found that the cutoff frequency decreased as a function of β (Fig. S1, Table S2).

To further understand the mechanism that fish used to modulate the low-frequency power of active movements as a function of reafferent gain, we segmented the swimming trajectories into “epochs”—continuous bouts of swimming in a single direction (Fig. S2). We examined how these epochs changed as a function of reafferent gain and lighting. There was a significant increase in the mean epoch duration in light in compared to dark (p_one-tail_<0.05, for 5 out of 6 individuals, Mann-Whitney-Wilcoxon test). We also observed that both the epoch distance (active movement between successive direction reversals) and epoch duration (time between successive direction reversals) increase significantly with refuge gain (*N*=6, Table S3). Thus, the overall mechanism for increasing RMS position (regulating RMS slip) is to swim farther for each active swimming epoch by increasing its duration rather than scaling up the overall swimming speed.

### Experimentally induced closed-loop filtering does not explain changes in active sensing

As a final control, we considered the possibility that the apparent change in active sensing could be explained solely by the change in closed-loop dynamics caused by our experimentally altered reafferent gain. In other words, what if the fish’s strategy were strictly open loop, but the resulting RMS fish motions were changing as an artifact of the (experimentally modulated) closed-loop dynamics?

To investigate this possible confound we modeled active sensing as resulting from an active probe signal, *a*(*t*), generated by the nervous system. We assumed that the fish’s task controller and swimming mechanics (task plant) are as described in [19,32]. The null hypothesis (open-loop active signal generation) would imply that the power spectral density of the probe signal *a*(*t*) should be invariant to the experimentally augmented feedback gain. In contrast, changes in the PSD of *a*(*t*) as a function of the feedback gain must arise from the feedback control in the animal. We illustrate these alternatives in Fig. 4 A: the active probe signal *a*(*t*) emerges from an “Active Signal Generator”. If the Active Signal Generator is operating in closed-loop, then it modifies the power spectrum of *a*(*t*) depending on feedback (dashed-line marked with “?” in Fig. 4 A).

**Figure 4:**
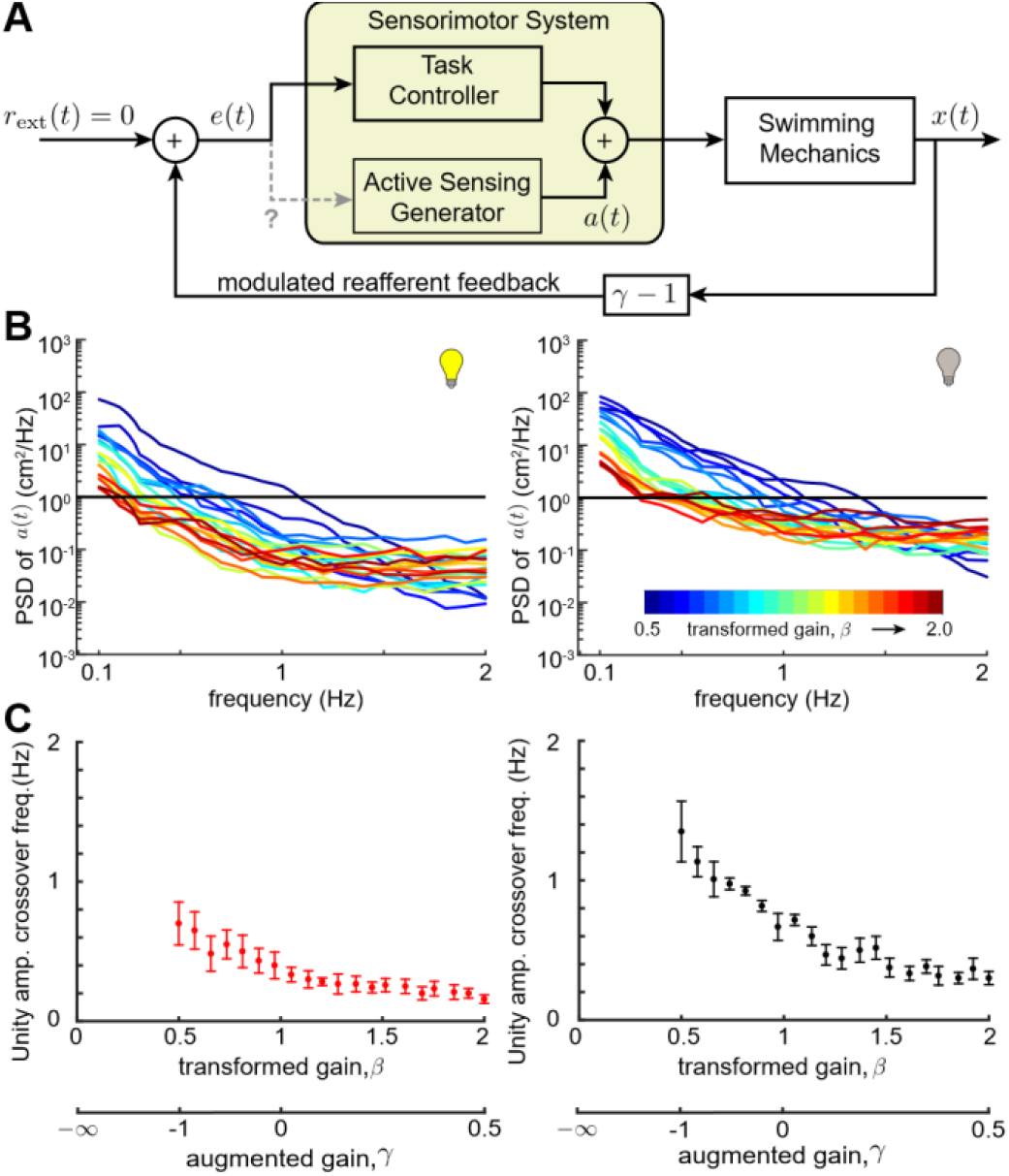
Open-vs. closed-loop control of active sensing. (A) Schematic of combined control of refuge tracking and active sensing. The active sensing signal is *a*(*t*). In the null hypothesis *a*(*t*) is open-loop, i.e. it is not modified in response to ongoing error feedback *e*(*t*). Rejecting the null model supports the hypothesis that the active sensing signal *a*(*t*) is modulated based on *e*(*t*), schematized by the gray dotted line (‘?’). The external input signal *r_ext_*(*t*) is zero in the present study. (B) Representative estimate of power spectra of *a*(*t*) for movements of an individual fish during ‘lights-on’ trials (left) and ‘lights-off’ trials (right). The solid black line corresponds to amplitude 1 cm^2^/Hz. We found that *a*(*t*) depends on β. (C) Combined data of unity-power cross over frequency of *a*(*t*) for all individuals (*N*=6) with mean ± SEM under different lighting conditions (red: ‘lights-on’, black: ‘lights-off’). See also Table S4.

For each gain, we estimated the PSD for *a*(*t*) using the following relationship:

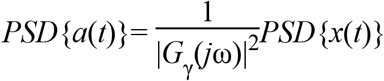

where *G*γ is the transfer function relating *a*(*t*) to *x*(*t*). We observed a substantial gain-dependent shift (two orders of magnitude) in the power spectrum of *a*(*t*) (Fig. 4 B,C, Table S4), supporting the hypothesis that fish dynamically regulate their active movements in relation to ongoing sensory feedback. These data reject the hypothesis that the closed-loop filtering properties of our augmented feedback system leads to the observed changes in active sensing behavior.

## Discussion

Previous experiments that modulated visual and electrosensory saliency [12] were used to establish the theory that fore-aft movements during tracking are for the purpose of active sensing. Similar forms of movement consistent with this theory have been described in another species of weakly electric fish [30]. The *a priori* hypothesis tested here, namely that the movements are regulated via reafferent feedback, was based on this theory of active sensing. Our experiments support this hypothesis and suggest that there are separate feedback control systems for active sensing and goal-directed behavior: one controller manages the flow of information and another task-level controller uses this information to achieve refuge tracking. We contend that the feedback regulation of active sensing is widespread across taxa and modalities, including whisking [4–6], touching [7,8], sniffing [9,10], and eye movements [11]. For example, in rats the timing of whisker protraction is locked to contact events, suggesting that whisking movements are under feedback control [24,33]. In humans, saccadic eye movements are triggered by low retinal slip [23], also suggesting that they are under feedback control.

Fish made adjustments in active-sensing movements to maintain sensory slip across manipulations of the gain of reafferent feedback both in the presence and absence of salient visual cues. The magnitude of sensory slip in the light was categorically less than in the dark, consistent with prior work [12]. In contrast, the relations between adjustments in conductivity—which affects electrosensory feedback but not vision or other modalities—and the magnitude of fore-aft movements [12] demonstrate that these movements are used almost exclusively for active sensing in the electrosensory domain. The lower set point for the magnitude of active-sensing movements in the light reflects a reliance on visual cues over electrosensory cues for tracking the position of the refuge. Indeed, fish dynamically lower their gain to electrosensory information for refuge tracking when salient visual cues are available [22].

Recent work has shown that ON and OFF cells in the hindbrain can give rise to bursts of activity that encode reversals in the direction of movement of looming/receding stimuli [34]. These sorts of bursts may also contribute to the detection of reversals of longitudinal movement [35]. In the midbrain, neurons encode velocity of longitudinally moving objects, so-called direction-selective neurons [36-40]; midbrain neurons are sensitive to specific ranges of temporal frequencies [41-43] and velocities of motion [40,44]. How do these computations relate to the control of active sensing? The maintenance of RMS slip across feedback gains was achieved by increasing the durations of epochs of sensory slip, while the numbers of reversals in the direction of sensory slip were not maintained across gains. This suggests that the control of active sensing is not tuned to regulate the stimulation of reversal-sensitive neurons. Instead, the changes in epoch duration but not the velocity of sensory slip suggest that the movements may be tuned to the temporal filtering properties of direction selective neurons.

Critically, neural circuits for active sensing are modulated in relation to task; active sensing movements are not conserved across electrosensory behaviors. For example the impulsive nature of prey capture movements [45] are categorically different than refuge tracking. Recent findings suggest potential substrates for task-dependent modulation of sensory processing via descending feedback pathways. For example, rather than relying solely on bottom-up computations, encoding of looming/receding objects is mediated by descending feedback from the midbrain [34]. Similarly, feedback rather than feed-forward information processing is involved in extracting electrosensory envelope information, a correlate of distance [46].

Active sensing movements may also ensure *observability* [47–49]—sufficient sensory information to enable estimation of the system state, the engineering analog of preventing perceptual fading. Indeed, beyond simply avoiding perceptual fading, feedback control of active movements may be used to optimize the flow of sensory information [15].

Closed-loop experimental approaches [6, 50] are crucial for disentangling complex interactions between sensing and control [51-53]. This is highlighted by the open-loop replay experiments: fish produced different output behavior based solely on whether the stimulus emerged from a closed-loop interaction or was replayed in open loop. This categorical shift in responses in open- and closed-loop may be a widespread feature of animal behavior [54, 55].

## Acknowledgments

Thanks to Kyle Yoshida and Ismail Uyanik for helping with data collection. This material is based upon work supported by a Complex Systems Scholar Award to NJC from the James McDonnell Foundation under Grant No. 112836, a Collaborative National Science Foundation Award to NJC and ESF under Grant Nos. 1557895 and 1557858, an NSF REU to LAA under Grant No. 1460674, and a Johns Hopkins Electrical Engineering graduate fellowship to DB.

## Author Contributions

Conceptualization: N.J.C., E.S.F., S.A.S., D.B., L.A.A.; Methodology: all authors; Software, D.B., L.A.A., B.P.V.; Formal Analysis: D.B. with S.A.S., L.A.A.; Investigation: D.B., L.A.A., and B.P.V.; Data Curation: D.B. Writing – Original Draft: D.B., L.A.A., E.S.F., and N.J.C.; Writing - Review & Editing: all authors; Visualization: D.B. with S.A.S., L.A.A., E.S.F., N.J.C.; Supervision: N.J.C.; Funding Acquisition: N.J.C. and E.S.F.

## STAR ⋆METHODS

### CONTACT FOR REAGENT AND RESOURCE SHARING

Further information and requests for data/protocols should be directed to and will be fulfilled by the Lead Contact, Noah J. Cowan.

### EXPERIMENTAL MODEL AND SUBJECT DETAILS

#### Subjects

Adult *Eigenmannia virescens* (10-15 cm in length) were obtained from commercial vendors and housed according to published guidelines [56]. The experimental tanks were maintained with a water temperature of ~27°C and conductivity in the range of 150-250 μS/cm. All experimental procedures were approved by the Johns Hopkins Animal Care and Use Committee and followed guidelines established by the National Research Council and the Society for Neuroscience.

### METHOD DETAILS

#### Experimental apparatus and procedure

The experimental apparatus was similar to that described in previous studies [20,22]. The test environment was a 17 gallon rectangular glass aquarium. The refuge was machined from a 111 mm (111.66±0.23) segment of 46.64×50.65 (46.64±0.33×50.65±0.10) mm gray rectangular PVC tubing. The bottom face of the tube was removed and a series of five rectangular windows (6 mm in width and spaced 19 mm apart) were machined into each side to provide visual and electrosensory cues. The refuge was suspended less than 0.5 cm above the floor of the tank by an acrylic mount attached to a linear stepper motor (STS-0620-R, HW W Technologies, Inc., Valencia, CA, USA).

Prior to trials, an individual fish was transferred to the testing tank and allowed to acclimate for 4-24 hr. During trials, the position of the fish inside the refuge was recorded using a video camera (pco.1200, PCO AG, Kelheim, Germany). Video was captured at 25 frames per second and the position of the fish was tracked in real time using custom vision software programmed in LabView (National Instruments, Austin, TX, USA). Altogether, this allowed an experimental paradigm where the refuge could be moved either with a gain directly proportional to the movements of the fish (experimentally closed loop) or with a specified trajectory (experimentally open loop).

The LabView-based video tracking algorithm employed template matching to determine the position of the fish in the video frames. For each fish, a custom image template was generated based on its appearance. Camera images were calibrated to determine the physical size of the area covered by each pixel in the plane that the fish swam so that we could estimate the physical location of the fish within the refuge. Given the requirement of low latency, we implemented an FPGA-based stepper motor controller in LabView that was directly controlled by the image-processing PC, sending it target positions as soon as the physical position of the fish was determined by the image-processing software. The stepper motor controller generated the fastest possible smooth trajectory to the target position and sent the corresponding pulse train to the motor driver. We controlled the camera frame rate using the same FPGA hardware along with the manual shutter feature of the camera. The capture frame rate was fixed at 25 frames per second and the image-capture-to-motor-control latency was less than one video frame time (<40 ms).

#### Closed-loop and open-loop experiments

Closed-loop experiments were 70 s in duration comprising three phases: Initiation (5 s duration), Cross-Fade (5 s) and Test (60 s duration). All of the data and analysis reported in these experiments are from the Test Phase.

In the Initation Phase, the refuge motion was a sinusoidal trajectory at 0.45 Hz, ramped over 5 s. As in previous studies [12,20], the gradual introduction of refuge motion reduces startle responses. Visual inspection of fish motion during this phase also provides a confirmation that the animal is attending to the tracking task (in this study, no trials were rejected based on visual inspection of the Initiation Phase). Pilot experiments were conducted without this Initiation Phase, and the animals were often startled at the beginning of the Test Phase. The parameters of the Initial Phase (duration, amplitude, frequency, ramp profile) were selected based on extensive prior experience with similar studies and fall well within the locomotor tracking performance (amplitude and bandwidth) of the fish. It is possible that details of the Initial Phase may affect details of the behavior during the Test Phase, so the same Initiation Phase parameters were used across all conditions to ensure consistency.

The Cross-Fade Phase provided a smooth transition from open-loop to closed-loop control of the refuge trajectory. During this phase, the amplitude of the open-loop sinusoidal stimulus was reduced to zero over a period of 5 s while the gain of the closed-loop component of refuge was concomitantly increased to its final test value for that experiment.

In the Test Phase of closed-loop experiments, the refuge motion was completely governed by the movement of the fish, as given in Eqn. 1. The choice of gains γ and its logic are explained later in *Details of closed-loop experiment*. In open-loop replay experiments, an entire 70 s refuge trajectory, previously recorded in a closed-loop trial, was presented to the animal. These trajectories included the refuge motion that resulted from the Initial, Cross-Fade, and Test Phases from a previously recorded trial. As in closed-loop trials, the analysis of open-loop replay trials was restricted to the final 60 s of the trial, corresponding to the Test Phase of the closed-loop experiment.

If the fish left the refuge or the real time tracking of the fish from the video feed was lost during a trial, the trial was terminated, and the data collected during that trial rejected. In such cases, the trial was repeated until completed successfully. This occurred in less than 10% of the trials, so no significant selection bias corrupted our observation.

#### Details of initiation and cross-fade phases

To smoothly initiate each trial, the refuge was first moved in a sinusoidal trajectory (0.45 Hz) which gradually ramped up in amplitude from 0 cm to 3 cm over the first 5 s of the trial (Initiation Phase). Over the next 5 s (Cross-Fade Phase), the refuge input smoothly transitioned from the sinusoidal trajectory to closed-loop control (Eqn. 1), in which the refuge motions were directly proportional to those of the fish by the specified gain constant, γ. Specifically, the refuge trajectory was defined by the following:

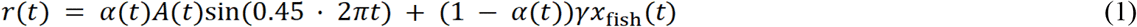

where

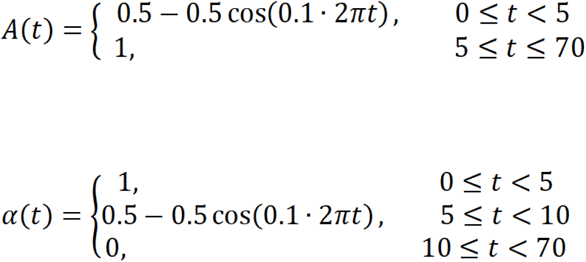

Here, *A*(*t*) specifies the smooth onset amplitude of the sinusoidal input which initiates the trial, while α(*t*) provides a “crossfade” parameter to smoothly transition from the initial sinusoidal input to closed-loop control.

#### Details of closed-loop experiment

Assuming constant sensory slip, *e_rms_* =*E*, we have

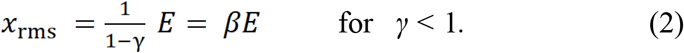

Note that a gain of γ=1 (β=∞) leads to a singularity in the closed-loop, whereby the fish is unable to change its relative position in the tube, ultimately causing the linear actuator to hit a travel limit. Gains of greater than 1 negates the sign of reafferent feedback, making it difficult for the fish to stabilize its position inside the refuge; such conditions, while potentially interesting, are beyond the scope of the present study.

To test the robust slip hypothesis, we selected a set of 20 proportional gain constants which were sampled along the real line between *γ*= −1 and *γ* = +0.5 such that values for β=(1-γ)^-1^ were uniformly spaced. This choice of gain spacing makes the comparison between theoretical expectations and experimental observations visually apparent: if slip is maintained constant by the fish, then we expected *x_RMS_* to be linear (with slope *E*) when plotted as a function of β.

We also tested the prediction that the RMS velocity was not affected by gain β. In this case, we predict that RMS of the sensory slip velocity, *e_v,rms_*, is related to β as follows:

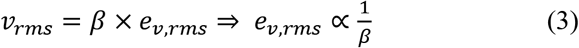

Thus we predict that *e_v,rms_* is in the shape of a hyperbole as function of β. The proportionality constant in Eqn. 3 can be obtained by fitting a constant to the RMS velocity data.

Six individuals (*N*= 6) were presented with closed-loop trials at each of 20 gain values. For each gain value, we performed trials with two complementary lighting conditions: ‘lights-on’ and ‘lights-off’. This gave a set of 40 unique trial conditions. We performed 3 replicate trials for each set of conditions, resulting in a total of 120 trials per individual. To reduce the possibility of learning or sequential ordering effects, we randomized the order of all trial conditions (gain and lighting). Each trial was separated by a rest period of 2-3 minutes. For any two successive trials with the same lighting condition, the opposite lighting condition was imposed during the rest period (for example, two consecutive ‘lights-on’ trials would be separated by a ‘lights-off’ rest interval).

#### Details of open-loop replay experiment

A separate open-loop replay experiment was designed to ensure that changes in behavior resulted from the coupling between the fish movement and the refuge movement, and not simply as a consequence of the refuge motion itself. Five individuals (N = 5) were presented with closed-loop trials using gain values of −1 and +0.22. For each gain value, the position of the refuge was recorded throughout three replicate closed-loop trials, giving three distinct refuge trajectories for playback. Each of these refuge trajectories was played back in five open-loop trials, in which the refuge motion pattern presented to the fish was the same trajectory as recorded in an earlier corresponding closed-loop trial. Additionally, five closed-loop trials were recorded for each gain value (these trials were not played back in open-loop) to offer further behavioral data for the closed-loop condition. The order of all trials was randomized for each individual, with the constraint that a closed-loop trial had to be completed (its trajectory recorded) before any of its five corresponding open-loop replay trials. This resulted in a total of 46 trials per fish (two gain values, each with three closed-loop trials recorded for playback, 15 open-loop playback trials, and five closed-loop trials for further comparison, one lighting condition: ‘lights-off’).

### Motion analysis

For each trial (*n*= 720 for closed-loop, *n* = 230 for open-loop), the digitized position of the fish for each frame was converted from raw pixel data to length units (centimeters), giving the longitudinal position of the fish as a time-varying signal over the period of a trial. The longitudinal position of the refuge for each trial was available directly in length units from the custom code which controlled the refuge (LabView, National Instruments, Austin, TX, USA). We calculated the RMS of the mean-subtracted longitudinal position of the fish, giving a single value per trial to represent the amount of motion of the fish. Occasional whole-body bending and transverse motion were not used in our analysis, thereby treating the refuge tracking behavior as a one-dimensional task [18.19]. From the mean-subtracted longitudinal position of the fish (*x*(*t*)), we calculated the velocity 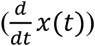, slip (*e*(*t*) = *r*(*t*) − *x*(*t*)), and slip-velocity 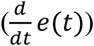to gain further insight regarding locomotor behavior. For the closed-loop experiment, we calculated RMS of each of these and averaged over each set of three replicate trials to characterize individual behavior across all trial conditions. For the open-loop experiment, we calculate the RMS of *x*(*t*) for each trial and compared the amount of movement between closed-loop trials and the corresponding open-loop playback trials. Please note since the signals are all mean subtracted, the RMS value represents the standard deviation of the probability distribution derived from the time series data.

### Analysis of open-loop replay data

For the statistical analysis of the open-loop replay experimental data we adopted the following simplification approach. For an individual, at a specific gain, we computed the ratio of RMS of each of the 5 open-loop replicates to the corresponding closed-loop data (movement/slip). We averaged these 5 ratios to get one value per closed-loop trajectory used in the open-loop replay experiment. Repeating the process for other two closed-loop trajectories we got three values and averaged these three values to get one representative value for an individual at a specific gain. Thus we ended up with total of two sets of 5 values corresponding to gain −1 and +0.22. The statistics reported in the main paper were performed on these datasets.

### Epoch analysis

Using a custom written MATLAB (MathWorks, Natick, MA, USA) script we detected the direction reversal points on time series active movement data. Based on this, we calculated the epoch distance (active movement between successive direction reversals) and epoch duration (time between successive direction reversals).

### Estimation of active probe signal from experimental data and system parameters

The original system and the plant transfer functions are taken as described in previous studies [17,28]. The transfer functions of task plant/swimming mechanics, *P*(*s*), and overall system, *H*(*s*), are given as follows:

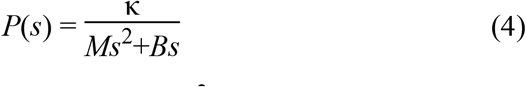

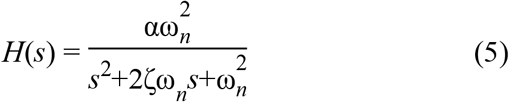

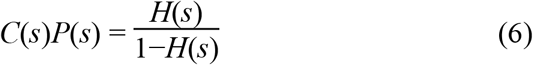

The parameter descriptions are from [19,21]:

DC gain, *α* = 1 [s^2^]; undamped natural frequency, *ωn* = 2*π* ×1.049 [rads^−1^]; damping coefficient, *ζ* = 0.56; mass constant, *M* = 2.8×10^−3^ [kg]; damping constant, *B* = 5.3×10^−3^ [kg s−1]; actuator gain, *κ* = 2.09×10^−3^ [kg s^−2^].

For the estimation of the active probe signal from the experimental data of the longitudinal position of the fish, *X*(*s*) we computed the transfer function, *G*γ(*s*) from *A*(*s*) to *X*(*s*) as follows:

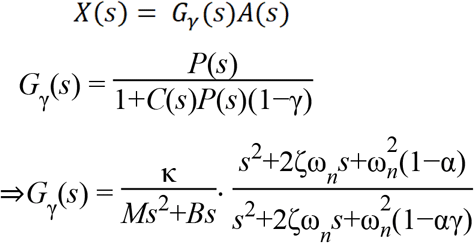

In the frequency domain,

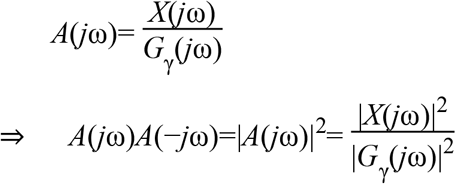

Under appropriate choice of window function, power spectral density (PSD) of active probe signal, *PSD*{*a*(*t*)}, and the longitudinal fish position, *PSD*{*x*(*t*)}, are related as follows:

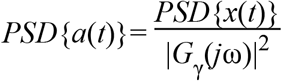

## QUANTIFICATION AND STATISTICAL ANALYSIS

All the statistical analysis were performed with custom codes written in MATLAB (MathWorks, Natick, MA, USA). The statistical tests used, as indicated in this manuscript, were as follows: one way ANOVA, Mann-Whitney Wilcoxon (MWW) test, Mann-Kendall test and Spearson’s rank correlation test. For all tests significance level was set to less than 0.05. The experimental data are provided as the mean plus or minus the standard error of the mean (μ±*SEM*).

## DATA AND SOFTWARE AVAILABILITY

An archived version of the datasets and the analysis code supporting this article will be made available through the Johns Hopkins University Data Archive with the following doi:10.7281/T1/DX9DL8 (https://archive.data.jhu.edu).

## Supplemental Video Files

**Video S1**. **Showing effect of negative gain**, *γ* = −1 **on fish movement. Related to Figure 1.** The blue line, aqua green line and the red line correspond to camera frame of reference, position of the fish, and position of the refuge, respectively. The top panel depicts the fish movement with respect to camera frame of reference (‘what we see’) whereas the middle panel shows the same with respect to the fish (i.e. fish position stabilized). Bottom panel shows refuge and fish trajectories. Enhancing the reafferent feedback (×2) results in less active movement.

**Video S2**. **Showing effect of positive gain, γ**= +0.22 **on fish movement. Related to Figure 1.** Same format as in Supplementary Movie 1. Suppressing the reafferent feedback (×0.78) results in more active movement.

**Figure S1.**
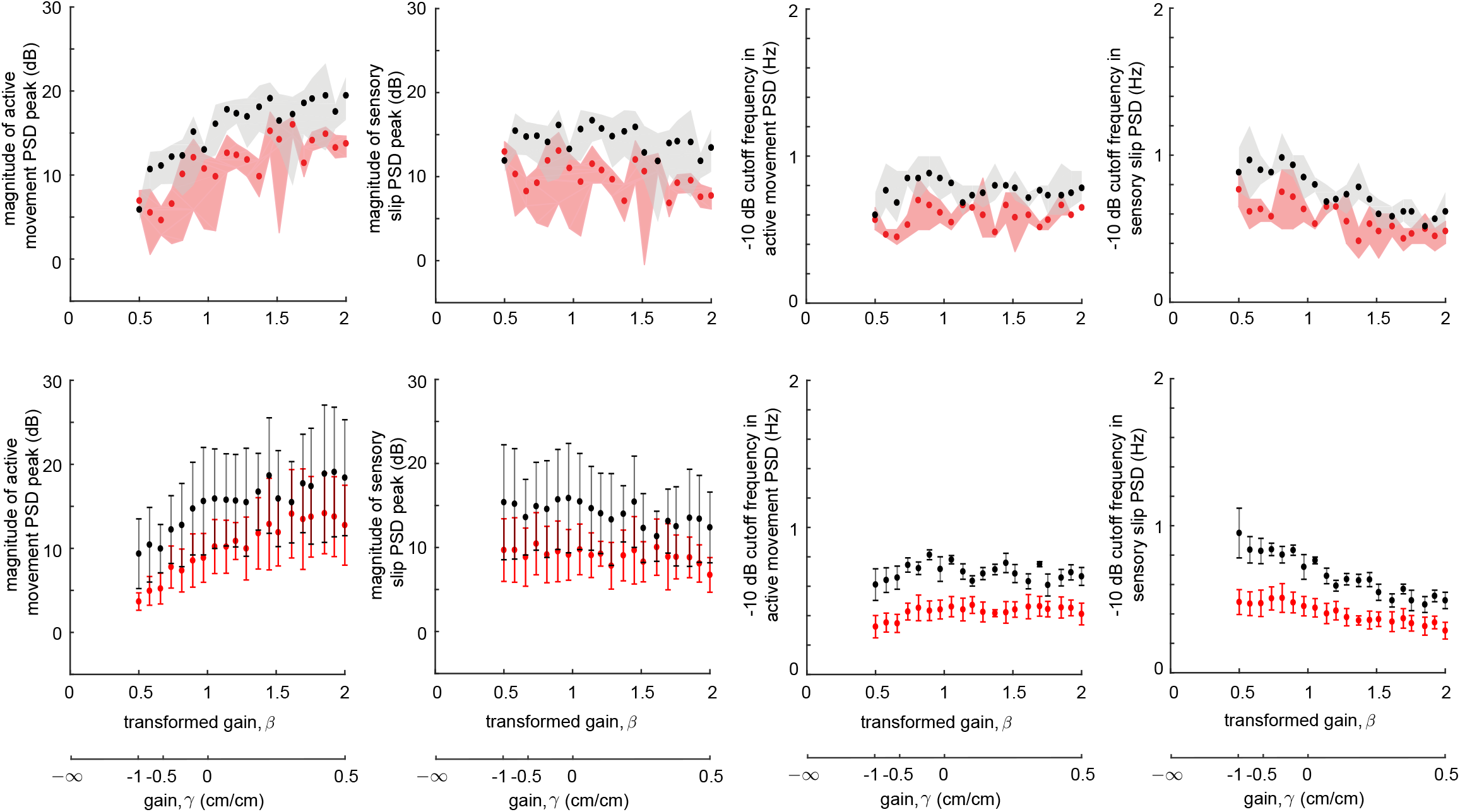
Variations of important features of power spectral density (PSD) of active movement and sensory slip across gains. Related to Figure 3. The top row presents representative data from one fish. Each marker represents respectively the peak amplitude of PSD of active movement (first column), peak amplitude of PSD of sensory slip (second column), 10 dB cutoff frequency of active movement PSD (third column) and 10 dB cutoff frequency of sensory slip PSD (fourth column) averaged across three replicates (black: ‘lights-off’, red: ‘lights-on’) at different gain values. The shaded regions denote the maximum and minimum experimental values at each gain. The bottom row represents combined data for all individuals (*N* = 6) with mean *±* SEM indicated.

**Figure S2.**
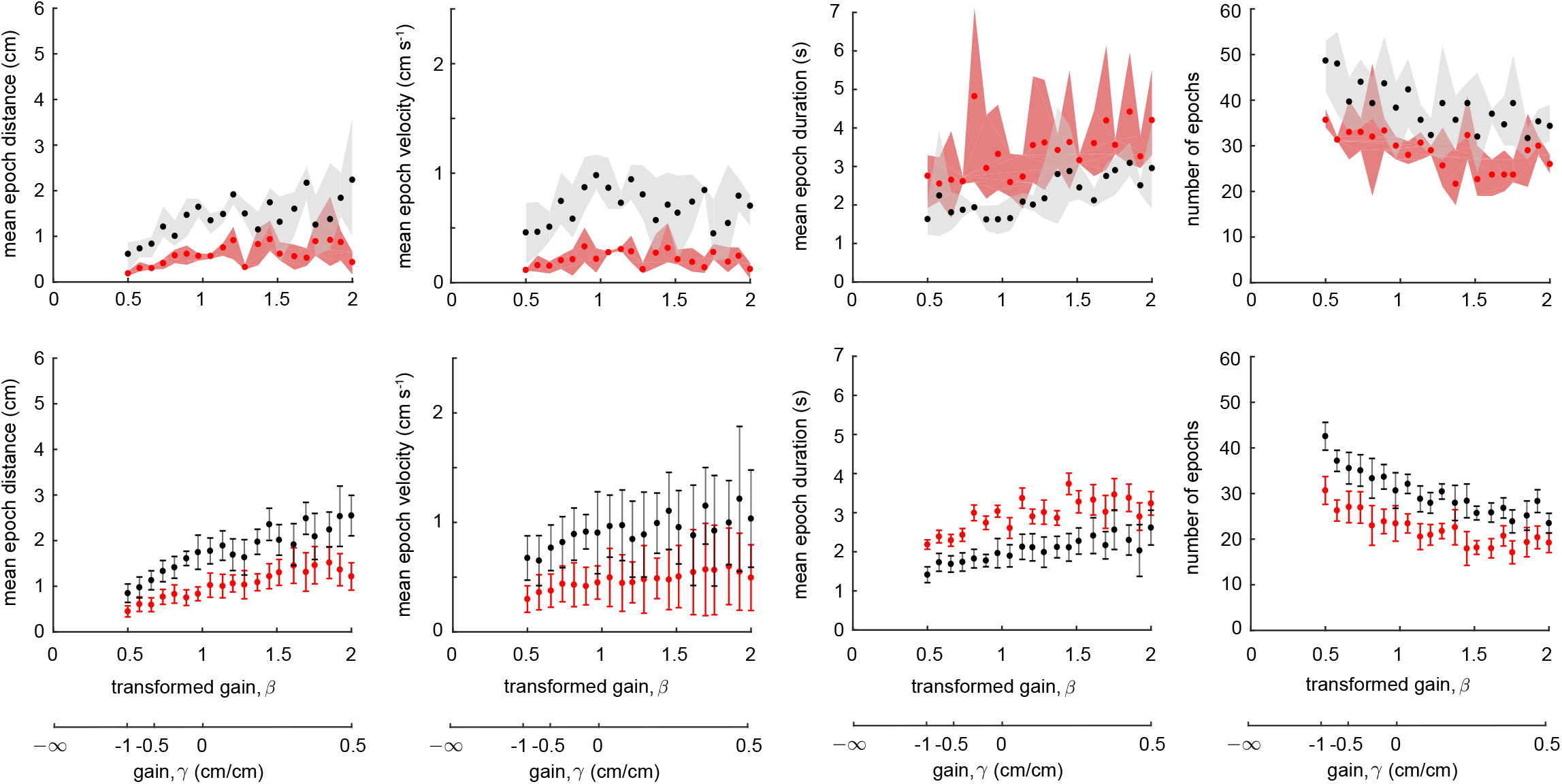
Epoch analysis. Related to Figure 3. The top row presents representative data from one fish. Each marker represents respectively the mean distance traversed by the fish between consecutive direction reversals, or *epochs* (first column), the mean epoch velocity (second column), the mean duration of the epochs (third column) and the number of epochs (fourth column) averaged across three replicates (black: ‘lights-off’, red: ‘lights-on’) at different gain values. The shaded regions denote the maximum and minimum experimental values at each gain. The bottom row represents combined data for all individuals (*N* = 6) with mean *±* SEM indicated.

**Table S1.**
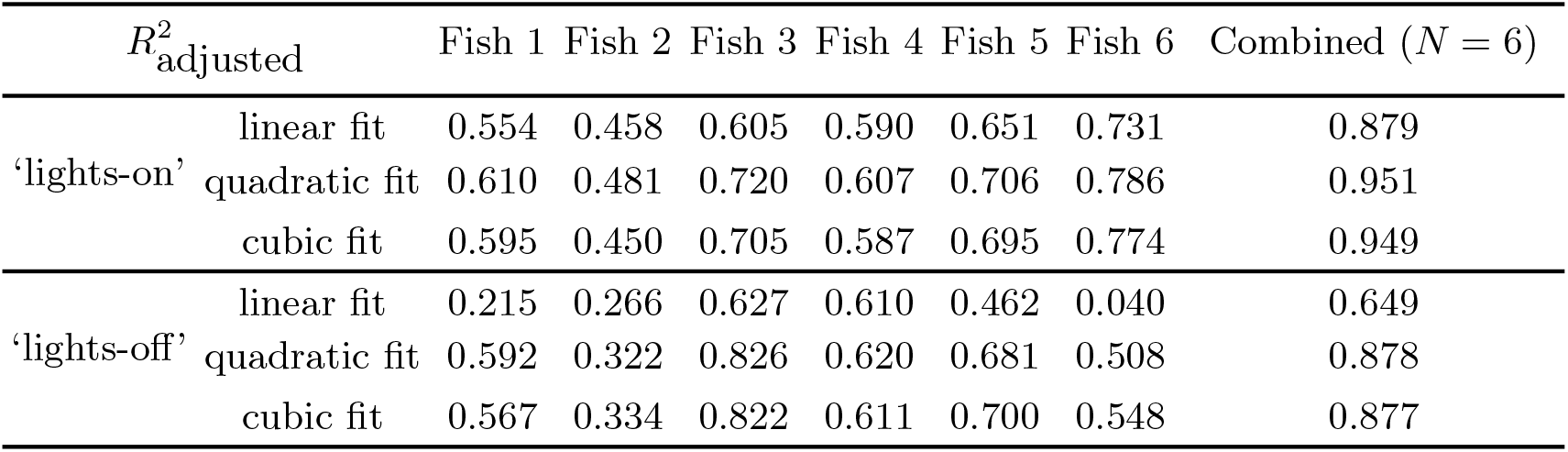
Adjusted *R^2^* values. Related to Figure 2. Adjusted *R*^2^ for linear, quadratic and cubic fit to the active movement data for each individual (*N* = 6) as well as to the combined data in both lighting conditions (‘lights-on’ and ‘lights-off’ trials).

**Table S2.** Mann-Kendall test and Sen’s slope estimator for detecting increasing/decreasing trend in PSD of active movement and sensory slip. Related to Figure 3. Low *p* value suggests existence of trend with respect to independent variable – transformed gain, *β*. Sen’s slope estimator is a robust unbiased estimator of the true slope of the linear regressor model fitted to the data.

| p-value | −10 dB cutoff frequency |  |  |  | PSD peak amplitude |  |  |  |
| --- | --- | --- | --- | --- | --- | --- | --- | --- |
|  | Active Movement |  | Sensory Slip |  | Active Movement |  | Sensory Slip |  |
|  | ‘lights-on’ | ‘lights-off’ | ‘lights-on’ | ‘lights-off’ | ‘lights-on’ | ‘lights-off’ | ‘lights-on’ | ‘lights-off’ |
| Fish 1 | 0.3649 | 0.6912 | 0.0458 | $2.65 \times 10^{-6}$ | $5.00 \times 10^{-5}$ | $3.17 \times 10^{-4}$ | 0.6732 | 0.6265 |
| Fish 2 | 0.2784 | 0.0103 | 0.03 | $3.95 \times 10^{-5}$ | $3.17 \times 10^{-4}$ | 0.0048 | 0.7212 | 0.2561 |
| Fish 3 | 0.5284 | 0.9463 | $3.49 \times 10^{-4}$ | $2.78 \times 10^{-6}$ | $5.17 \times 10^{-4}$ | $1.34 \times 10^{-6}$ | 0.556 | 0.2561 |
| Fish 4 | 0.0628 | 0.0626 | 0.0013 | $4.86 \times 10^{-5}$ | $1.60 \times 10^{-5}$ | $8.32 \times 10^{-4}$ | 0.3468 | 0.381 |
| Fish 5 | 0.8915 | $7.00 \times 10^{-4}$ | $4.61 \times 10^{-5}$ | $2.51 \times 10^{-6}$ | $2.85 \times 10^{-5}$ | $1.91 \times 10^{-4}$ | 0.7212 | 0.0297 |
| Fish 6 | 0.3883 | 0.4915 | 0.0111 | 0.0019 | $4.93 \times 10^{-7}$ | 0.0021 | 0.6265 | 0.1441 |
| sen’s slope | −10 dB cutoff frequency |  |  |  | PSD peak amplitude |  |  |  |
|  | Active Movement |  | Sensory Slip |  | Active Movement |  | Sensory Slip |  |
|  | ‘lights-on’ | ‘lights-off’ | ‘lights-on’ | ‘lights-off’ | ‘lights-on’ | ‘lights-off’ | ‘lights-on’ | ‘lights-off’ |
| Fish 1 | 0 | 0 | −0.090 | −0.449 | 6.127 | 23.028 | −0.745 | −1.697 |
| Fish 2 | 0.058 | −0.146 | −0.1483 | −0.378 | 15.191 | 32.228 | −1.635 | −13.076 |
| Fish 3 | 0 | 0 | −0.157 | −0.318 | 14.757 | 56.780 | −4.236 | −5.548 |
| Fish 4 | 0.130 | 0.066 | −0.191 | −0.210 | 13.269 | 40.367 | 0.932 | −6.648 |
| Fish 5 | 0 | −0.148 | −0.247 | −0.488 | 27.781 | 65.039 | −1.745 | −14.400 |
| Fish 6 | 0.017 | −0.079 | −0.091 | −0.339 | 9.708 | 12.058 | 0.343 | −3.821 |

**Table S3.**
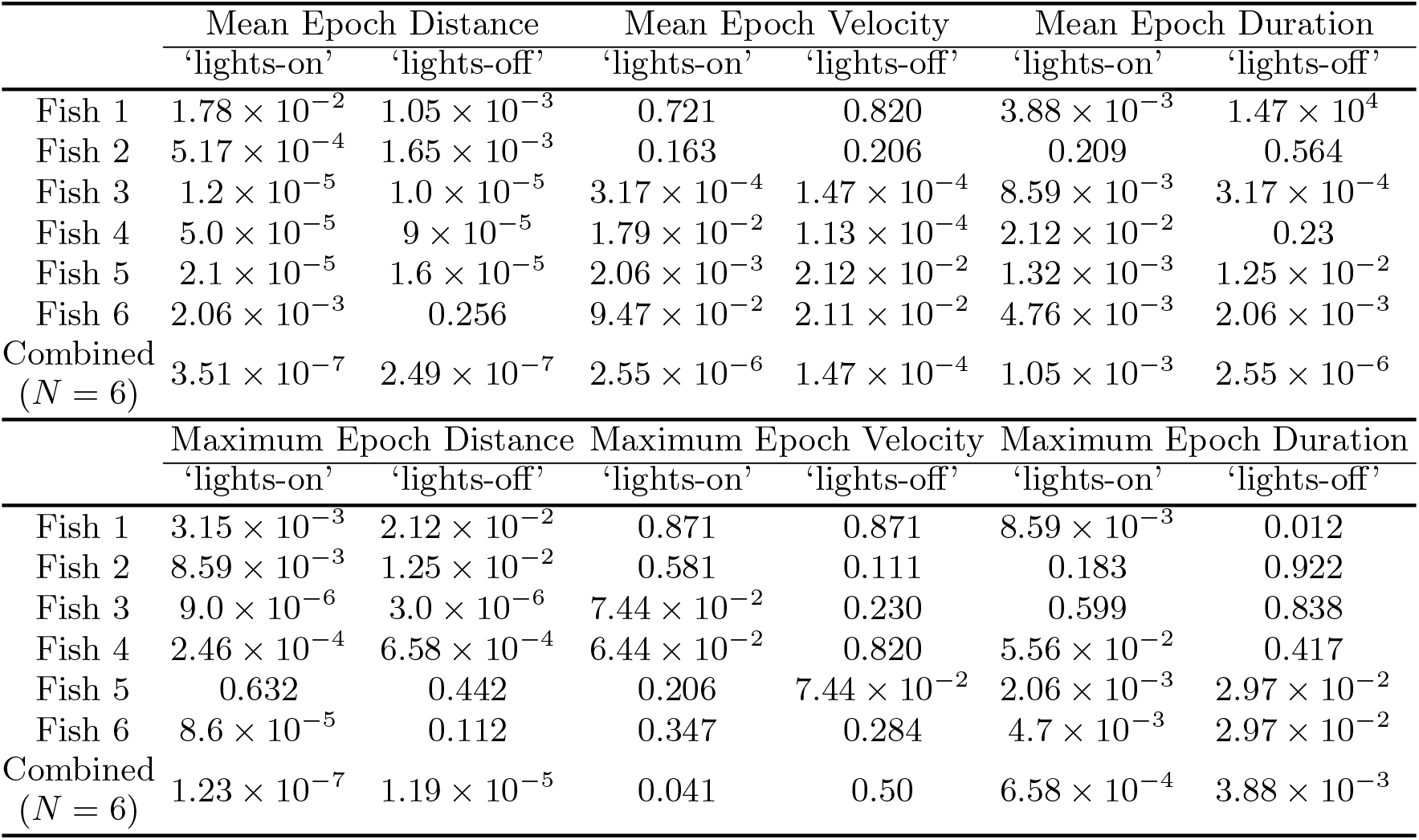
Mann-Kendall test for detecting trend in mean and maximum epoch parameters – distance, velocity and duration. Related to Figure 3. Low *p* value suggests existence of trend with respect to independent variable – transformed gain, *β.*

**Table S4.** Mann-Kendall test for detecting trend in PSD of active probe signal, *a*(*t*). Related to Figure 4. Low *p* value suggests existence of trend with respect to independent variable – transformed gain, *β.*

|  | Unity amplitude crossover frequency |  |  |  |  |  |
| --- | --- | --- | --- | --- | --- | --- |
|  | Fish 1 | Fish 2 | Fish 3 | Fish 4 | Fish 5 | Fish 6 |
| 'lights-on' | $0.2476 \times 10^{-3}$ | $0.1149 \times 10^{-3}$ | $0.0074 \times 10^{-3}$ | $0.0057 \times 10^{-3}$ | $0.0136 \times 10^{-3}$ | $0.0698 \times 10^{-3}$ |
| 'lights-off' | $0.0008 \times 10^{-3}$ | $0.2927 \times 10^{-3}$ | $0.0001 \times 10^{-3}$ | $0.0145 \times 10^{-3}$ | $0.0014 \times 10^{-3}$ | $0.0271 \times 10^{-3}$ |

